# A chromosome-level genome assembly of the chilli thrips *Scirtothrips dorsalis* Hood (Thysanoptera: Thripidae)

**DOI:** 10.64898/2026.10.08.757792

**Authors:** Allan Busuulwa, Tracy Liesenfelt, Andrew J. Mongue, Sriyanka Lahiri

**Author notes:** Corresponding Author: Allan Busuulwa.

## Abstract

Thrips are known for being pests of many agricultural crops, as well as vectors of plant viruses. Yet despite their economic importance, genomic resources for this group remain limited, and no chromosome-level genome has been available for the genus *Scirtothrips*. One such species is the chilli thrips, *Scirtothrips dorsalis* Hood (Thysanoptera: Thripidae). This invasive and highly polyphagous pest is established across much of the world and transmits several tospoviruses. Genomic resources would provide a new means to understand the traits that make thrips difficult to manage. Here, we present a chromosome-level genome assembly for *S. dorsalis*. The assembly is 266.5 Mb, with 96.9% of the sequence anchored in 16 chromosome-scale scaffolds. It contains 20,557 genes, and repetitive elements account for 28.8% of the sequence, which is comparable to those of other sequenced Thripidae. A species tree based on 3,019 single copy orthologs placed *S. dorsalis* as sister to *Scirtothrips hansoni,* with *Scirtothrips* most closely related to *Frankliniella, Megalurothrips,* and *Odontothrips*. The mitochondrial genome is a single circular molecule of 15,207 bp in which the sequence of the published mini circle is integrated, in contrast to the bipartite mitogenome previously reported for the same cryptic species.

## Introduction

*Scirtothrips dorsalis* Hood (Thysanoptera: Thripidae) (Figure 1) is currently a global pest of many economically important crops, particularly ornamentals, vegetables, and fruits (Kumar et al. 2013; Dickey, Kumar, Hoddle, et al. 2015; Busuulwa et al. 2026). However, there is significant contradiction regarding the origin of *S. dorsalis.* Mound and Palmer (1981) reported the species to be native to the region extending from Pakistan in the west to Japan, Solomon Islands and Queensland to the east. Seal et al. (2006) reported *S. dorsalis* to have originated from South Asia while Hoddle et al. (2008) reported it to be native to Australasia and South Africa. Kumar et al. (2013) reported it to be native to the Indian subcontinent while in 2013 CABI reported it to be native to the Indo-Pacific region especially in Bangladesh, Myanmar, Pakistan, Sri Lanka, Taiwan, Thailand, Papua New Guinea and the Solomon Islands, and as an introduced species in South Africa.

**Figure 1.**
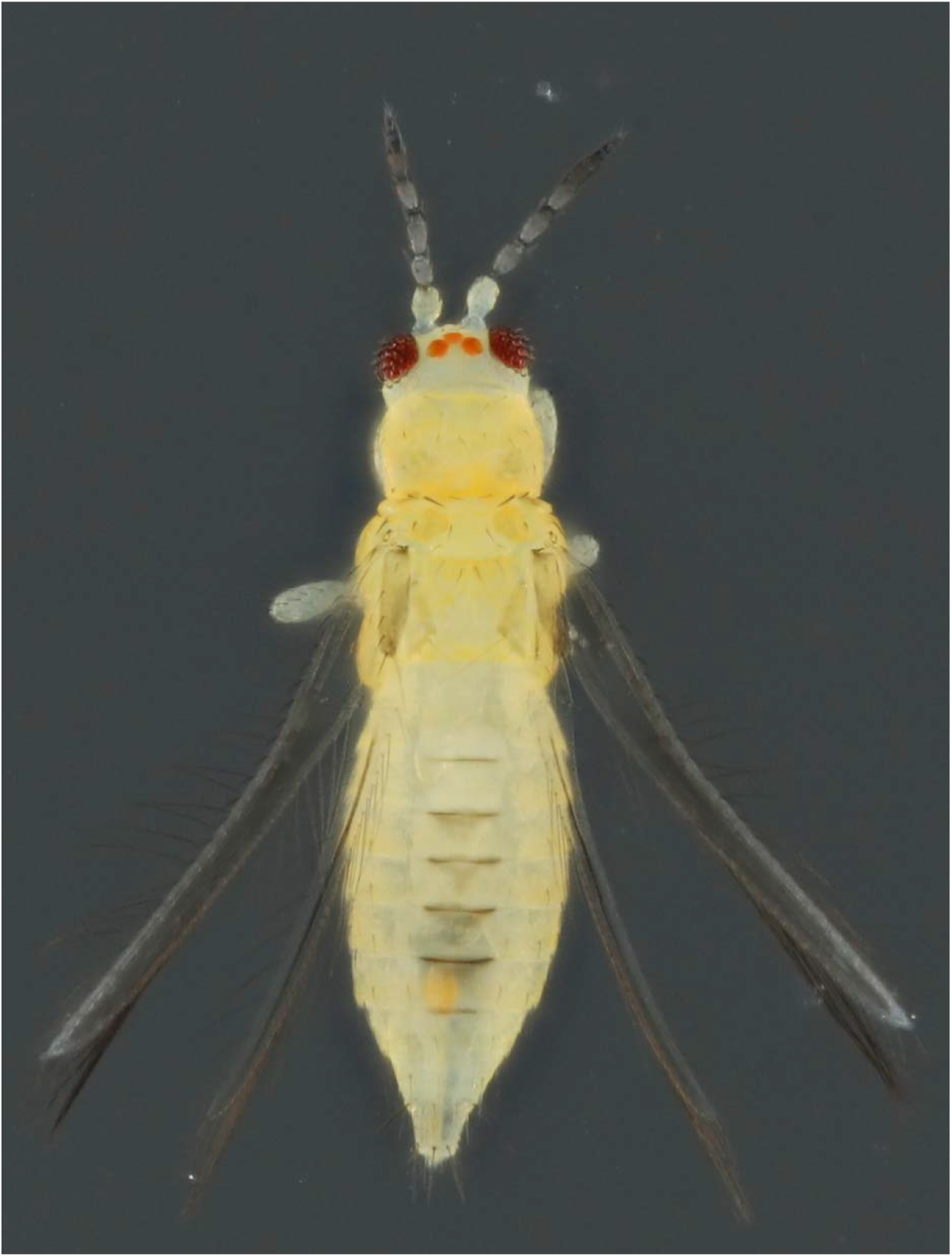
Adult female of *Scirtothrips dorsalis* Hood. Photo by AB.

Therefore, based on where the reported native ranges overlap, the most likely origin is South Asia, specifically the Indian subcontinent, from where the species may have spread east through Southeast Asia into the western Pacific and Australia. This is supported by the original description of *S. dorsalis* from specimens collected on chilli, *Capsicum annuum* L. (Solanaceae) and castor, *Ricinus communis* L. (Euphorbiaceae) in India (Hood 1919). Much of the controversy regarding the origin of *S. dorsalis* can be explained by the fact that *S. dorsalis* is not a single species but rather a species complex with each having its own native range. Dickey et al. (2015) recognized 9 cryptic species (i.e. South Asia 1 and 2; East Asia 1, 2, 3, and 4; and Australia 1, 2, and 3) and 2 morphologically distinguishable species (i.e. *Scirtothrips oligochaetus* and *Scirtothrips* aff. *dorsalis*). Iftikhar et al. (2016) later identified 11 mitochondrial cytochrome C oxidase subunit 1 gene (COI) lineages corresponding to separate barcode index numbers (BINs) on the Barcode of Life Data System (BOLD), thereby suggesting that this complex might contain up to 11 cryptic species, although this remains to be confirmed by nuclear markers. Additionally, the same study found 7 *S. dorsalis* haplotypes in Pakistan, with 3 shared with India, further supporting the origin of *S. dorsalis* to be the Indian subcontinent.

Nonetheless, the range of this pest has expanded rapidly to include parts of the Middle East, Africa and the Americas (Kumar et al. 2013). Two cryptic species have been reported in the USA. South Asia 1 is well established in the southern states, including Florida and Texas, while East Asia 1, which is native to northern Japan, has been found in the Northeast and appears able to overwinter in regions with freezing winters (Kumar et al. 2023). In Florida, *S. dorsalis* (South Asia 1) is currently a major pest of strawberry, *Fragaria x ananassa* Duchesne (Rosaceae) (Busuulwa, Revynthi, et al. 2024a, 2024b; Busuulwa, Riley, et al. 2024; Lahiri et al. 2024) and blueberry, *Vaccinium corymbosum* L. (Ericaceae) (Lahiri et al. 2025), where it feeds on young leaves causing curling and scarring on fruits causing bronzing.

Management of *S. dorsalis* is challenging for several reasons. Larvae and adults tend to shelter in unopened leaves, buds and other young growth, where they are hard to detect and are protected from contact insecticides and predators (Kaur and Lahiri 2022). In strawberry, for example, females lay their eggs inside the leaf tissue, where they are protected from predators (Adhikary, Smith, Whitaker, et al. 2025), while the remaining life stages also occur on the leaves, often in sheltered sites This cycle is completed quickly under warm conditions. At around 30 °C, *S. dorsalis* develops from egg to adult in roughly 14 days (Tatara 1994). As a result, it can complete up to 8 generations per year in temperate regions and as many as 18 in the warmer southern United States, allowing populations to build up rapidly (Nietschke et al. 2008). Its broad host range also allows it to survive on weeds and other alternative hosts when preferred hosts are absent, therefore permitting it to survive all year-round (Lahiri and Perry 2025). As a result, management in many crops has relied heavily on repeated insecticide applications, which has led to reduced susceptibility of *S. dorsalis* to several commonly used insecticides (Kaur et al. 2023). This limitation has resulted in increased interest in alternative management strategies, especially biological control (Busuulwa, Revynthi, et al. 2024a; Lahiri et al. 2024; Adhikary, Smith, and Lahiri 2025), and has further emphasize how little is known about the genetic basis of traits such as insecticide resistance and host use in this species.

Despite the economic importance of thrips, genomic resources for the group currently remain limited. Chromosomal genome assemblies are currently present for a few pest species, most of which are in the Thripidae family. These currently include *Thrips palmi* Karny (Guo et al. 2020), *Stenchaetothrips biformis* (Bagnall) (Hu et al. 2023), *Megalurothrips usitatus* (Bagnall) (Ma et al. 2023; Zhang et al. 2023) *Frankliniella intonsa* (Trybom) (Song, Wang, et al. 2024), *Frankliniella occidentalis* (Pergande) (Song, Cao, et al. 2024), *Odontothrips loti* (Haliday) (Yingning et al. 2024), *Thrips tabaci* Lindeman (Gao et al. 2024) and *Pseudodendrothrips mori* (Niwa) (Guan et al. 2026). Within *Scirtothrips* genus, the only currently available genome is a highly fragmented assembly of *Scirtothrips hansoni* Mound & Hoddle (360,059 scaffolds, scaffold N50 3.6 kb), while none has been published for *S. dorsalis*. This gap therefore limits research into the genetic basis of traits that make *S. dorsalis* difficult to manage and also leaves its relationship to other thrips untested with genome scale data. Furthermore, a chromosomal-level genome reference would help refine the status of the cryptic species within the complex, which are currently distinguished mainly by mitochondrial and a small number of nuclear markers.

The present study therefore aimed to (i) assemble and annotate a chromosome-level genome of *S. dorsalis* from a Florida colony using PacBio HiFi and Illumina Hi-C sequencing, (ii) assemble and characterize its mitochondrial genome, including the arrangement of the region corresponding to the published mini-circle, and (iii) determine its phylogenetic position among the available thrips genomes.

## Materials and Methods

### Scirtothrips dorsalis sample collection

The DNA sequence was obtained from a mixture of approximately 180 larvae and adult *S. dorsalis* of both sexes, taken from a laboratory colony that had been originally collected from strawberry fields locally and kept at the University of Florida-Gulf Coast Research and Education Center, Wimauma, FL, U.S.A., since 2016. Over that period the colony was maintained continuously on cotton *Gossypium hirsutum* L. (Malvales: Malvaceae) in a growth room set at 25 ± 1 °C, 65 ± 5% RH relative humidity and a 16:8 light: dark photoperiod and had never been exposed to insecticides or fungicides. Furthermore, no individuals were introduced from wild populations after the colony was established, therefore by the time of sample collection, the colony had become a distinct inbred line, which allowed pooling DNA from different individuals without introducing substantial genetic variation. The collected *S. dorsalis* individuals were frozen at −80 °C for approximately five minutes, after which DNA extraction began immediately.

### DNA extraction and sequencing

*Scirtothrips dorsalis* tissue samples collected for DNA extraction were homogenized using a Dounce homogenizer directly into the lysis buffer provided in an Omniprep DNA extraction kit for high-quality genomic DNA (G-Biosciences, St. Louis, MO, USA). The standard Omniprep base kit protocol was used with minor adjustments as described in Liesenfelt et al. 2025. These adjustments had been successfully used to extract high weight molecular DNA from both single individual insects (Liesenfelt et al. 2025) and pooled samples of multiple individuals (Mongue et al. 2026). In brief, the adjustments included lysing the tissue samples overnight for 15-16 hours on an incubator (Fisher brand, mini heat block, Waltham, MA, USA) at 56 °C, adding 2 μL of 20 mg/mL Mussel glycogen (Thermo Scientific, Waltham, MA, USA) to increase DNA yield during the DNA precipitation step, and allowing DNA to precipitate for 1 hour in a −20 °C freezer.

### Quality control and sequencing

The quantity and quality of the DNA were assessed using a Qubit 4 Fluorometer (Thermo Fisher Scientific, Waltham, MA, USA) with a Qubit dsDNA 1X Broad Sensitivity Assay Kit and a NanoDrop spectrophotometer (Thermo Fisher Scientific, Waltham, MA, USA). The extracted DNA was subsequently sent to the Arizona Genomics Institute (Tucson, AZ, USA) for Pacific Biosciences (PacBio) HiFi sequencing on a Revio system. Additionally, approximately 50 mg of tissue (> 15,000 *S. dorsalis*) was sent to the same facility for Illumina Hi-C library preparation and sequencing. Three samples, each containing approximately 500 individuals (mixed stages of both sexes), were sent to GENEWIZ (South Plainfield, NJ, USA) for paired-end RNA sequencing.

### Mitochondrial genome and species confirmation

To confirm the identity of the sequenced material, the HiFi reads were first used to assemble the mitochondrial genome. Since *S. dorsalis* is a complex of cryptic species with high variability in their mitochondrial architecture (Dickey, Kumar, Hoddle, et al. 2015; Kumar et al. 2023), it was necessary to manually select the mitogenome reference beforehand instead of accepting one from an auto search. To select the best mitogenome to act as a reference, all available *S. dorsalis* mitochondrial sequences between 800 bp to 20 kb available in GenBank (n = 50) were retrieved. HiFi reads were then aligned to each mitogenome separately using minimap2 v2.28 (Li 2018) by leveraging the **map-hifi** flag. The strategy of aligning each reference independently was chosen because mapping all of them together at once would assign each read to whichever reference fitted it best while leaving the others without reads no matter the similarity.

Mapping results indicated that the HiFi reads matched most closely (95% identity) with chromosome 1 of KM349827.1, the *S. dorsalis* cryptic species delineated as South Asia 1. Therefore, KM349827.1 was chosen as the reference, and all HiFi reads that aligned to it were used as the mitochondrial set. The mitochondrial genome was then assembled from this set of reads using MitoHiFi v3.2 (Uliano-Silva et al. 2023) under the invertebrate mitochondrial genetic code. Since mitochondria are present in many copies per cell, the set of reads used in the assembly was deeply covered, at approximately 975-fold the expected size of the *S. dorsalis* mitogenome based on published mitogenomes (Dickey, Kumar, Morgan, et al. 2015). This was far more than what hifiasm (Cheng et al. 2021), the assembler MitoHiFi runs internally is currently designed for. Therefore, the reads were subsampled using seqkit v2.13.0 (Shen et al. 2016) to 50,100 and 200-fold and each subsample assembled independently. The three assemblies were then compared to each other, and the one chosen was verified by mapping the mitochondrial reads back to it using minimap2 and examining the read depth in 100 bp windows using SAMtools v1.20 (Danecek et al. 2021), followed by aligning the assembly against itself to detect internal repeats. Gene annotation of the assembled mitochondrial genome was done using both MitoFinder v1.4.0 (Allio et al. 2020) and MITOS2 v2.1.0 (Donath et al. 2019), after which the two annotations were reconciled. In cases where the two annotations disagreed, the sequence in question was used to settle the conflict, by comparison with the reference and by examining the anticodons of the transfer RNA genes. Gene order was then compared directly with that of the reference rather than by whole mitochondrial genome alignment, since rearrangement in thrips is extensive enough to make such alignment uninformative (Tyagi et al. 2020).

Since the published *S. dorsalis* South Asia 1 mitogenome is bipartite (*i.e.* the mitogenome is split across two separate circular chromosomes), with *nad6* and *trnC* on a 921 bp chromosome of their own (Dickey, Kumar, Morgan, et al. 2015), the reads were examined to establish whether the sequences assembled here came from one or both chromosomes. To do this, the stretch of the assembly corresponding to the published mini circle was first located. It ran from a copy of *trnS1*, through the control region, *trnC* and *nad6*, to a second copy of *trnS1*. The mitochondrial reads were then mapped back onto the assembly so that this stretch and its junctions could be examined read by read.

First, reads crossing each of the segment’s two junctions were counted, requiring that at least 500 bp of sequence on either side be found nowhere else in the mitogenome and that the alignment contain no gaps, so that a read could not have aligned there by chance. These counts were compared with junctions elsewhere in the mitogenome. Secondly, read depth across the segment was compared with the depth of the rest of the genome. Third, the number of copies of the segment appearing in individual reads was counted. The identity of this segment to the published chromosome 1 and mini circle sequences was measured using Smith-Waterman alignment with EMBOSS water v6.6.0 (Rice et al. 2000), both gene by gene and across the segments as a whole. Additionally, both circular references were doubled beforehand, such that a query spanning the origin would still align. Lastly, COI sequence was extracted from the annotated mitogenome using a custom bash script and queried against the BOLD Systems (Ratnasingham and Hebert 2007).

### Whole genome size estimation

Jellyfish v.2.3.0 (Marçais and Kingsford 2011) was used to estimate the expected genome size of *S. dorsalis* using the raw HiFi reads. K-mer frequencies were generated with **jellyfish count -m 21 -s 3G -C** flags. The resulting histogram was analyzed using GenomeScope2 v2.0 (Ranallo-Benavidez et al. 2020) to estimate the coverage depth of the homozygous regions and the size of the genome.

### Genome assembly

#### Contig assembly from raw reads

Hifiasm v0.25.0 (Cheng et al. 2021) was used to assemble the primary genome with the **-l 3** parameter to purge all types of haplotigs. The remaining haplotigs and overlaps were subsequently removed with purge_dups v1.0.1 (D. Guan et al. 2020), using coverage cutoffs calculated from HiFi reads mapped back to the primary assembly using minimap2. The assembly was then assessed for completeness using the Insecta ortholog dataset (insecta_odb12) in BUSCO v6.0.0 (Tegenfeldt et al. 2025). The BUSCO assessment of this assembly returned 98.5% complete but 91.7% duplicated, with only 211 of 3,114 genes present in a single copy. Duplication at this level is not explained by unpurged haplotigs, and suggested redundancy in the input reads rather than in the assembly. Subsequent screening of the raw HiFi reads with seqkit showed that the reads had 1,752,330 records corresponding to 876,165 unique read names, meaning that every read was present twice. Therefore, the raw reads were deduplicated using **seqkit rmdup -n** and the primary assembly repeated as previously described.

#### Decontamination of the PacBio primary assembly

After the primary assembly and purging of duplicated haplotigs, BlobTools v4.3.5 (Challis et al. 2020) was used to search the assembly for potential contaminants and endosymbionts. The purged assembly was then split into 1 FASTA per contig using seqkit to allow for each to be searched independently against the NCBI core_nt database using megablast (BLAST v2.15.0, Altschul et al. 1990), with an e-value cutoff of 1e-25, and a maximum of 10 target sequences per query as recommended in the BlobTools documentation. Given that roughly a third of the assembly was repetitive, a culling limit of 5 was used to remove hits contained within better scoring hits. The resulting hits were then combined into a single hits file using a custom bash script.

To obtain coverage information, the deduplicated HiFi reads were mapped back to the primary assembly using minimap2, and the alignments sorted and indexed using SAMtools. Afterwards, the hit file, the coverage from the alignment, the BUSCO full table from the earlier Insecta_odb12 run and the NCBI taxonomy databases were combined into a BlobDir with *S. dorsalis* (NCBI taxid 163899) set as the target taxon. Blob and snail plots were generated to visualize the distribution of contigs by GC content, coverage and taxonomic assignment.

To remove contaminating sequences, the taxonomic assignment of each contig was tested. Contigs assigned to Streptophyta and Arthropoda were extracted separately using SAMtools followed by assessment with BUSCO against Insescta_odb12. This was done to determine whether the contigs that had been identified as plants carried any insect single-copy orthologues. Contigs assigned to Arthropoda or no hits were retained and written to a cleaned assembly while all others assigned to Streptophyta, Chordata or Mycoplasmatota were discarded. No hit contigs were kept because *S. dorsalis* currently had no close relatives that were well represented in the nucleotide databases, meaning that an absence of hits could not be used as evidence for contamination. The cleaned assembly was then accessed for completeness using BUSCO against Insescta_odb12 and for quality using mercury v1.3 (Rhie et al. 2020).

#### Hi-C scaffolding

To resolve the assembly to chromosomal level, the decontaminated primary assembly was scaffolded using Hi-C sequencing data. Hi-C reads were mapped to the assembly following the <u>Arima pipeline</u> (Arima Doc A160156 v03). In brief, each read pair was aligned separately to the primary assembly using BWA-MEM v0.7.17 (Li and Durbin 2009). Next, the alignments generated were passed through the Arima 5′ filter to retain only the 5′ most alignments of reads spanning a ligation junction, followed by pairing the two ends and filtering them to a minimum mapping quality of 10. Read groups were then added, PCR duplicates removed using <u>Picard</u> v3.2.0, and the resulting alignments sorted and indexed using SAMtools.

The resulting alignments were used to create an initial scaffolded assembly using YaHS v1.2.2 . (Zhou et al. 2023), with the minimum mapping quality (**-q**) of 10 to match the filter applied during mapping and the contig error correction (**--no-contig-ec**) disabled. Error correction was disabled because YaHS breaks contigs where Hi-C contact coverage is sparse, and this library was not dense enough to distinguish a genuine misjoin from thin coverage, whereas the contigs themselves derived from 44× HiFi data and had already been assessed against the read depth distribution. Thereafter, a JBAT file of the initial scaffolded assembly was generated using Juicer Tools v1.22.01 (Durand, Shamim, et al. 2016) and then visualized as a Hi-C contact map using Juicebox (Durand, Robinson, et al. 2016) to manually curate misassembles. The curated scaffolded assembly was then assessed for completeness and quality using BUSCO and Mercury respectively. BUSCO assessment of the scaffolded assembly showed a high duplication rate, so a second round of haplotig purging was done using purge_dups, followed by a further BlobTools screen to identify and remove any remaining contaminants as described above.

### Repeat annotation and masking

Transposable elements (TE) were annotated, and the genome soft-masked using Earl Grey v7.2.6 (Baril et al. 2024). Earl Grey is a complex automated pipeline that combines several tools into a single workflow to curate and annotate transposable elements in a genome. In brief, it first identifies elements de novo using RepeatModeler2, followed by refining each consensus sequence through an iterative process of BLAST searching, extraction, alignment and trimming, thereby extending truncated models towards full length. The refined library built is then passed onto RepeatMasker to annotate and mask the genome, after which RepeatCraft is used to defragment and de-overlap the resulting annotations so that a single element previously broken into pieces is reported as one feature rather than several.

In this run, RepeatModeler2 v2.0.8 (Flynn et al. 2020) and <u>RepeatMasker v4.2.3</u> were used together with the Dfam 3.9 library, a reference database of transposable element families curated from a wide range of organisms. HELIANO was enabled in the run in order to detect Helitons, which standard tools currently identify poorly. Since *S. dorsalis* had not been previously screened for repeats, no reference library was supplied to the initial masking step to allow for elements to be identified de novo. This was done to prevent novel TE families from being forced into ones previously described. Additionally, consensus clustering was disabled during the run because clustering collapses chimeric elements and closely related subfamilies, thereby distorting divergence estimates. The soft-masked assembly produced by this step was subsequently used as the input for gene prediction.

### Gene prediction

Two independent annotations, one from protein homologues and the other from RNA sequencing evidence were performed and then combined. For the protein homology annotation, a custom Thysanoptera protein database containing 10 thrips species was assembled from published annotations (Table 1). Four sets were downloaded from NCBI and the remaining downloaded from Figshare where the majority had been deposited. Protein sequences for *O. loti*, were extracted from its deposited genome and annotation using gffread v0.12.7 (Pertea and Pertea 2020), while all other sets were used as deposited. After combining the protein sets, all duplicated protein sequences were removed from using seqkit.

**Table 1.** Thysanoptera protein sets used as homology evidence for gene prediction.

| Species | Source | Accession or DOI | Proteins |
| --- | --- | --- | --- |
| *Frankliniella fusca* | NCBI | GCA_029379305.1 | 26,340 |
| *Frankliniella intonsa* | Figshare | 10.6084/m9.figshare.24174591.v5 | 14,109 |
| *Frankliniella occidentalis* | Figshare | 10.6084/m9.figshare.24968679.v1 | 16,312 |
| *Megalurothrips usitatus* | NCBI | GCA_026979955.1 | 12,782 |
| *Odontothrips loti* | Figshare | 10.6084/m9.figshare.24865023.v2 | 20,128 |
| *Pseudodendrothrips mori* | Figshare | 10.6084/m9.figshare.30256126.v2 | 13,429 |
| *Stenchaetothrips biformis* | Figshare | 10.6084/m9.figshare.23708619.v1 | 17,167 |
| *Thrips hawaiiensis* | Figshare | 10.6084/m9.figshare.26125162 | 19,024 |
| *Thrips palmi* | NCBI | GCF_012932325.1 | 27,217 |
| *Thrips tabaci* | Figshare | 10.6084/m9.figshare.24408181.v1 | 17,816 |
| **Total** |  |  | **184,324** |

Gene models were predicted with GALBA v1.0.11 (Brůna et al. 2023) on the soft-masked assembly, using the custom Thysanoptera database as the sole source of evidence. In brief, GALBA aligns proteins to the genome using miniprot (Li 2023) and then uses the alignments to train AUGUSTUS (Hoff and Stanke 2019) and extrinsic evidence during prediction. GALBA was chosen for this task mainly because Thysanoptera is represented by only a small number of annotated genomes, making high-quality proteins only available from a few closely related species rather than from a large clade database.

In the second annotation that leveraged RNA sequences, reads were first trimmed using fastp v0.23.4 (Chen 2025), then aligned to the soft-masked genome using with HISAT2 v2.2.3(Kim et al. 2019) using the **--dta** option to produce alignments suited for transcript assembly which were then sorted and indexed using SAMtools. Subsequent gene prediction was carried out using <u>BRAKER4</u> v0.5.0-beta, implemented in Snakemake (Köster and Rahmann 2018), in ET mode, which reproduces the BRAKER1 pipeline (Hoff et al. 2019). Within this pipeline, spliced alignments were converted to introns with bam2hints from AUGUSTUS v3.5.0, while strand orientation was determined and spurious introns removed using **filterIntronsFindStrand.pl** based on recognized splice site dinucleotides. GeneMark-ET (Lomsadze et al. 2014) was then trained on these hits and successively used to produce an initial gene set that was filtered to only retain genes with evidence support.

DIAMOND v2.0.15 (Buchfink et al. 2021) was subsequently used to remove redundant sequences. To reduce over-representation of single-exon genes, the remaining training genes were down-sampled using a Poisson distribution with a lambda of 2. The resulting set was then split into a training set and a validation set of 300 genes. AUGUSTUS species specific parameters were trained using etraining and further refined with **optimize_augustus.pl**. Predictions from AUGUSTUS and GeneMark-ET were merged using TSEBRA v1.1.2.5 (Gabriel et al. 2021), and the merged set compared against the AUGUSTUS and GeneMark sets using **best_by_compleasm** (Brůna et al. 2024) to run compleasm (Huang and Li 2023) in protein mode to retain sets that contained the fewest missing BUSCOs. The final set was filtered for internal stop codons and the CDS coordinates normalized. UTR features were added by decorating the gene models with StringTie2 (Kovaka et al. 2019) transcript assemblies.

Annotations based on the protein dataset and RNA-seq were subsequently combined using TSEBRA using the **keep_ab_initio** configuration in order to retain transcripts that the extrinsic evidence did not support rather than discarding them. This configuration was chosen because many of the protein-derived models lacked RNA-seq intron support, which the default thresholds required, and were therefore being discarded.

### Functional annotation

The merged set was converted from GTF to GFF3 with **agat_convert_sp_gxf2gxf.pl** from AGAT v1.4.2 (Jacques Dainat et al. 2026) followed by single transcript per gene selection using **agat_sp_keep_longest_isoform.pl**. Protein sequences were extracted from the longest-isoform models using gffread against the soft-masked assembly, after, stop-codon characters and lowercase residuals arising from masked regions were removed. The protein sets were subsequently assessed for completeness using BUSCO against Insecta_odb12 in protein mode. Protein domains, families and Gene Ontology terms were assigned utilizing InterProScan v5.75-106.0 (Jones et al. 2014) by searching Pfam, SMART, SUPERFAMILY, CDD and PANTHER with GO term and pathway lookup enabled.

Functional assignments based on orthology were obtained using eggNOG-mapper v2.1.15 (Cantalapiedra et al. 2021) by searching against eggNOG v5 database (Huerta-Cepas et al. 2019) while restricting search to Insecta (taxonomic scope 6656) and retaining only non-electronically inferred Go terms. The results were combined into a single annotation using fuannotate v.1.8.17 (Palmer and Stajich 2020), which additionally searched UniProtKB/Swiss-Prot, MEROPS, dbCAN and the Insecta BUSCO set, assigned gene names and product descriptions, and applied NCBI-format locus tags to every gene model.

Furthermore, DIAMOND was used to access homology to curated and non-curated protein databases, particularly UniProtKB/Swiss-Prot and the arthropod subset of UniProtKB/TrEMBL, using **`--very-sensitive`** mode with an e-value cutoff of 1e-5 while requiring that alignments cover at least 50% of both query and subject. It is, however, important to mention that the two databases were searched separately so as to independently report hit rates from each database.

#### Non-coding RNA annotation

Non-coding RNAs were annotated against the final assembly, starting with the transfer RNAs that were predicted utilizing tRNAscan-SE v2.0.12 (Chan et al. 2021) in eukaryotic mode. Ribosomal RNAs and other structured RNAs were predicted by scanning the genome against the Rfam database (Ontiveros-Palacios et al. 2025) using Infernal v1.1.4 (Nawrocki and Eddy 2013) with Rfam’s curated gathering thresholds, while microRNAs were predicted using MirMachine v0.3.0.4 (Umu et al. 2023) against the Arthropoda node of MirGeneDB. Since Rfam identifies tRNA and microRNA families alongside other non-coding RNAs, hits to those families were removed to avoid duplicating the tRNAscan-SE and MirMachine predictions at the same loci. The remaining Rfam features were merged with the tRNA, microRNA predictions and the functionally annotated protein-coding gene models into a single release annotation, sorted and written in GFF3 format with sequence-region headers matching the assembly.

#### Chromosomal architecture

Chromosomal scale scaffolds were first divided into 100 kb windows using bedtools v2.30.0 (Quinlan and Hall 2010). Afterwards, gene density, repeat content, GC content and non-coding RNA density were calculated for each window. The density of genes and non-coding RNA were obtained by counting features overlapping each window utilizing **bedtools intersect** on the functionally annotated gene models and non-coding RNA predictions. After merging the overlapping repeat intervals, repeat content was calculated as the fraction of each window covered by repeat annotations obtained from Earl Grey. This was done separately for all interspersed repeats, unclassified elements, and classified elements, which pooled DNA transposons, LTR and LINE elements, rolling-circle elements and satellites. It is important to mention that simple repeats and low-complexity sequences were not included in the interspersed repeat total. GC content was calculated for each window with **bedtools nuc** and then averaged across 5 neighboring windows, spanning 500 kb. The tracks were plotted in R v4.6.1 (R Core Team 2026) using circlize v0.4.16 (Gu et al. 2014), and non-coding RNA density was square root transformed so that windows with few features remained visible.

Satellite repeat unites were directly reconstructed from HiFi reads using SRF (Y. Zhang et al. 2023). SRF was selected as a tool of choice because it leverages frequently occurring k-mers to build repeat units, allowing the recovery of satellite arrays that are frequently collapsed during assembly. In brief, k-mers of 151 bp were counted using KMC V3.2.1(Kokot et al. 2017) and only those present at least 10 times the average read coverage were assembled into repeat units with SRF. In order to locate the arrays, each repeat unit was joined end to end into a longer sequence using **srfutils.js**. The assembly was then aligned to these sequences using minimap2 followed by the merging of overlapping alignments. To separate terminal arrays from internal ones, the position of each array was expressed as a proportion of its chromosome length. Telomeric repeats were identified utilizing seqkit by searching the assembly for 4 tandem copies of the canonical insect telomeric motif TTAGG (Y. Zhou et al. 2022). Matches within 20 kb of a chromosome end were counted as terminal telomeric arrays.

#### Phylogenetic placement

To determine the phylogenetic position of *S. dorsalis* among Thripidae, a species tree was inferred from single copy orthologs of all publicly available thrips genomes. *Scirtothrips hansoni* (GCA_059711435.1) was included as the only other genome available at the time for the genus, although the assembly was highly fragmented (448.1 Mb in 360,059 scaffolds with a scaffold N50 of 3.6 kb). Given that Hemiptera is a sister group to Thysanoptera (Tyagi et al. 2020; Johnson et al. 2018), the pea aphid *Acyrthosiphon pisum* (Hemiptera: Aphididae) was used as the closest outgroup while the body louse *Pediculus humanus* (Psocodea: Pediculidae) was added as a more distant outgroup, since Psocodea falls outside both orders (Johnson et al. 2018).

Single copy orthologues were identified in each genome using BUSCO against Insecta_odb12, the same dataset used to assess assembly and annotation completeness. BUSCO was used in place of an all-against-all approach such as OrthoFinder because it operates directly on genome assemblies, therefore allowing the highly fragmented *S. hansoni* and *F. occidentalis* assemblies to be used without being annotated first. The resulting genes recovered as complete and single copy in at least 80% of taxa were retained for further analysis. Protein sequences from each of the identified genes were aligned using MAFFT v7.520 (Katoh and Standley 2013) and the alignments trimmed with trimAl v1.4.1 (Capella-Gutiérrez et al. 2009) using the **-automated1** flag, with alignments shorter than 50 amino acids after trimming discarded. Amino acid sequences were used instead of nucleotides because Thysanoptera and Hemiptera diverged approximately 407 million years ago (Johnson et al. 2018), and synonymous sites are likely to be saturated over this timescale.

The species tree was then inferred using two approaches in order to avoid relying on a single set of assumptions. First, a concatenated maximum likelihood tree was built with IQ-TREE v2.4.0 (Minh et al. 2020), with each gene as a separate partition and models selected using ModelFinder (Kalyaanamoorthy et al. 2017). Branch support was assessed with 1,000 ultrafast bootstraps (Hoang et al. 2018) and 1,000 SH-aLRT replicates. A coalescent tree was also inferred with ASTRAL-III v5.7.7 (C. Zhang et al. 2018) from individual gene trees to account for incomplete lineage sorting. Both trees were inferred unrooted and then rooted on *P. humanus* for display.

## Results and Discussion

A total of 23.49 Gb of PacBio HiFi data containing 1,752,330 reads was received from the sequencing core (Table 2). Given that the library was found to contain every read twice, the reads were deduplicated, leaving 876,165 reads totaling 11.75 Gb and representing approximately 44× coverage of the assembled genome, with a read N50 of 15,210 bp and a mean length of 13,406 bp. Hi-C sequencing produced 48,221,874 read pairs totaling 13.98 Gb, while RNA sequencing of three samples produced 106,289,203 read pairs totaling 31.89 Gb.

**Table 2.** Sequencing data generated for *Scirtothrips dorsalis*.

| Library | Platform | Reads | Bases (Gb) | Mean (bp) | N50 (bp) | Max (bp) | Q30 (%) | GC (%) |
| --- | --- | --- | --- | --- | --- | --- | --- | --- |
| HiFi, as received | PacBio Revio | 1,752,330 | 23.49 | 13,406 | 15,210 | 54,593 | 92.95 | 52.78 |
| HiFi, deduplicated | PacBio Revio | 876,165 | 11.75 | 13,406 | 15,210 | 54,593 | 92.95 | 52.78 |
| Hi-C R1 | Illumina | 48,221,874 | 6.99 | 145 | — | 145 | 94.07 | 55.55 |
| Hi-C R2 | Illumina | 48,221,874 | 6.99 | 145 | — | 145 | 93.50 | 55.61 |
| RNA-001 R1 | Illumina | 36,145,503 | 5.42 | 150 | — | 150 | 94.79 | 52.03 |
| RNA-001 R2 | Illumina | 36,145,503 | 5.42 | 150 | — | 150 | 94.09 | 52.52 |
| RNA-002 R1 | Illumina | 35,523,770 | 5.33 | 150 | — | 150 | 94.56 | 54.13 |
| RNA-002 R2 | Illumina | 35,523,770 | 5.33 | 150 | — | 150 | 94.11 | 54.73 |
| RNA-003 R1 | Illumina | 34,619,930 | 5.19 | 150 | — | 150 | 94.22 | 56.01 |
| RNA-003 R2 | Illumina | 34,619,930 | 5.19 | 150 | — | 150 | 93.61 | 56.39 |

### Mitochondrial genome

As previously mentioned, the HiFi reads matched most closely with Chromosome 1 of the *S. dorsalis* cryptic species delineated as South Asia 1 (KM349827.1) at 95% identity. This was further corroborated by the BOLD system query which returned *S. dorsalis* for all 100 retrieved records, with 22 matching at 100% identity and no indels. Additionally, the query was assigned to BIN BOLD:AAC9748, which accounted for all matches above 98.2% identity. Likewise, some records were assigned to 2 more *S. dorsalis* BINs, BOLD:AAC9749 and BOLD:ACQ4218, matching at 96.5 to 96.8% respectively. This is consistent with the recognized division of *S. dorsalis* into cryptic lineages (Kumar et al. 2023; Dickey, Kumar, Hoddle, et al. 2015). These results therefore identified the sequenced colony as South Asia 1, the lineage that has been established in Florida since 2005 (Kumar et al. 2023). In addition, the reads also matched the 921 bp South Asia 1 mini circle (KM349828.1) at 98.9% identity, thus indicating the presence of both parts of the published *S. dorsalis* bipartite mitochondrial genome in the colony. Subsampling of the reads produced 3 assemblies in which the assembly from the 50-fold coverage was 15,207 bp while that of both the 100- and 200-fold coverage were 16,128 bp and contained a 922 bp tandem duplication with only 583-fold coverage compared to the 975-fold across the rest of the mitochondrial genome. This was an indication of assembly error and as a result, the 50-fold assembly was retained.

The assembled mitogenome contains the full set of all the 37 expected genes in an insect mitochondrion, comprising of 13 protein codding, 2 ribosomal and 22 transfer RNA genes (Figure 2). It is important to note that both MitoFinder and MITOS did not successfully detect the transfer RNA gene *trnV*. It was instead identified by its sequence, which was identical to the annotated copy in KM349827.1 which is consistent with the reported reduced structure of *trnV* in thrips (Tyagi et al. 2020)

**Figure 2.**
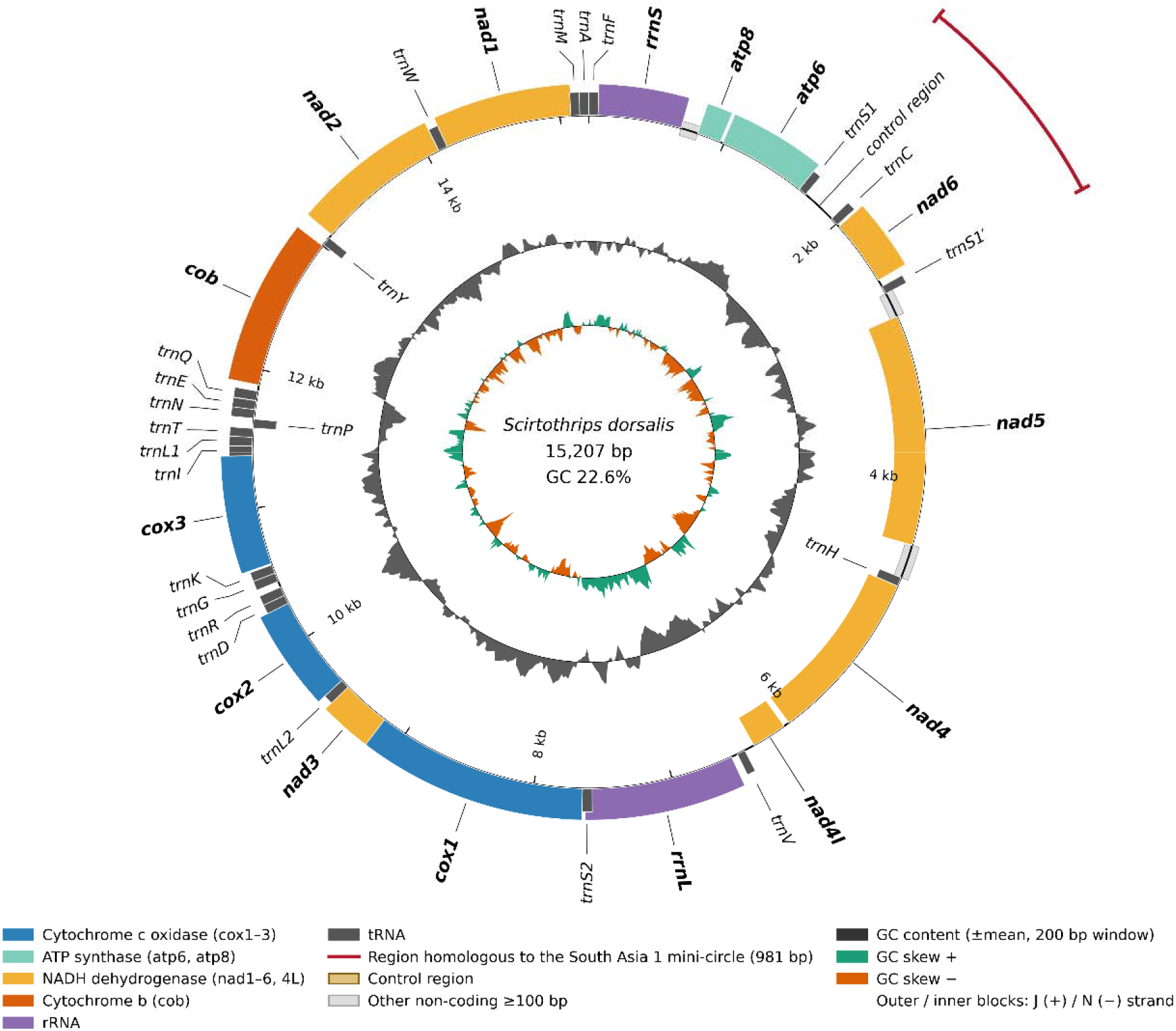
Mitochondrial genome of *Scirtothrips dorsalis* (South Asia 1). The mitogenome is a single circular genome of size 15,207 bp containing all the expected 37 genes in an insect mitochondrion. Genes on the outer ring are encoded on the J (+) strand while those on the inner ring are encoded on the N (-) strand and are further colored based on their functional group. Th read arc shows the region that matches the published South Asia 1 mini circle. Here, it lay between *atp6* and *nad5* and is flanked by two similar copies of *trnS1* (*trnS1* and *trnS1*′ ) and is found within the main mitogenome.

Gene order matched that of chromosome 1 of the South Asia 1 *S. dorsalis* cryptic species with one exception involving the 2 genes that define the mini circle. In the published mitogenome of South Asia 1, *nad6* and *trnC* sit on a separate 921 bp chromosome together with the duplicated copies of the control region and *trnS1* and not in the large chromosome (Dickey, Kumar, Morgan, et al. 2015), while here, the whole mini circle sits within the large chromosome between *atp6* and *nad5*. On the other hand, a 981 bp segment at this position matched the published mini circle at 99.3% identity along its full length with the control region at 98.1% identity and *trnS1*, *trnC* and *nad6* each at 100%. Additionally, the same segment matched chromosome 1 of South Asia 1 at only 49.8% mainly because that chromosome entirely lacks *nad6* and *trnC*. Furthermore, the segment is flanked by two identical 59 bp copies of *trnS1*, while the original location of *nad6* and *trnC*, between *nad4l* and *trnV*, is an empty 75 bp gap, as in the reference.

The three tests used to determine whether the segment formed part of the main mitochondrial genome, or was a separate chromosome joined to it during assembly, showed that it genuinely is part of the main chromosome. The segment’s two junctions with the flanking sequences were spanned by 872 and 792 reads, which was comparable to the 757 to 1,012 reads meeting the same criteria at other junctions in the mitogenome. Read depth across the segment was also 1.01 times the genome average, which was consistent with the segment being present in the same copy number as the rest of the genome. No reads contained more than two copies of the segment, and those that had two copies spanned the origin of the circular genome, as expected of reads averaging 13 kb on a 15.2 kb circle. A separate 921 bp mini circle sequenced with reads of this length would instead appear as multiple tandem copies within individual reads.

Dickey, Kumar, Morgan, et al. (2015) described this mini circle as the first multipartite mitochondrial genome in Thysanoptera and attributed it to a recent duplication of the control region. The arrangement found here, with the mini circle intact within the large chromosome, flanked by two identical copies of *trnS1* and carrying its own control region, is consistent with insertion of the mini circle into the main chromosome. However, because the original position of *nad6* and *trnC* is empty, this arrangement does not represent the state before fragmentation. The present data cannot show which arrangement came first given that the duplicated copies are too similar to tell apart. They do however suggest that one population of South Asia 1 carries this region as part of a single chromosome, whereas the published isolate carried it as two separate chromosomes. Therefore, it appears that mitochondrial genome structures vary between populations of the same cryptic species and the split originally described by Dickey, Kumar, Morgan, et al. (2015) is not fixed across *S. dorsalis* populations of South Asia 1

### Genome size estimation

K-mer analysis of the deduplicated reads estimated a haploid genome size of approximately 335.5 Mb, of which 112.0 Mb was estimated to be repetitive and 223.4 Mb, or 66.6% of the total, unique (Figure 3). The k-mer spectrum was single-peaked at a coverage of 32.4, and the model returned 100% homozygosity, as expected from a highly inbred colony.

**Figure 3.**
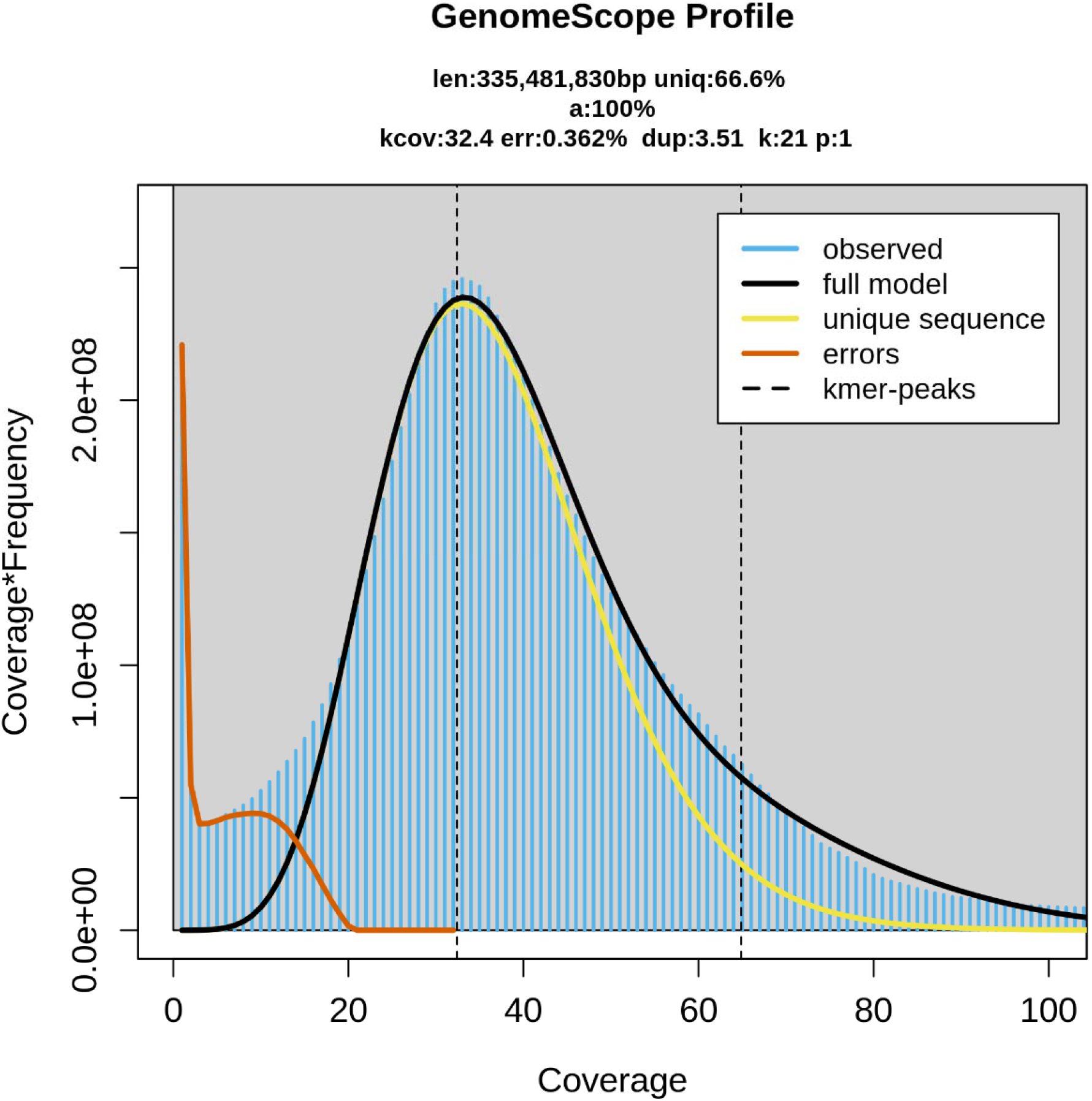
GenomeScope2 profile of 21-mers from the deduplicated *Scirtothrips dorsalis* HiFi reads on linear coverage scales. Observed k-mer counts are in blue, the fitted full model in black, the unique-sequence component in yellow and the error component in orange. Dashed lines mark the fitted k-mer peak at 32.4-fold and its second multiple.

### Genome assembly and Scaffolding

The initial assembly of the deduplicated HiFi reads produced 1,627 contigs that spanned 504.75 Mb with a contig N50 of 2.17 Mb, which was higher than the estimated genome size. BUSCO assessment of this assembly gave C:98.4% [S:51.9%, D:46.5%], F:0.4%, M:1.3%, indicating that approximately half of the single copy orthologues were present twice in the assembly. Subsequent purging of haplotigs reduced the assembly to 233 contigs spanning 342.19 Mb, with a duplication of 12.3% and a contig N50 of 4.23 Mb.

Contamination screening removed 61.06 Mb and 103 contigs, leaving 130 contigs spanning 281.13 Mb. Scaffolding the assembly with Hi-C data resolved the remaining contigs into 54 scaffolds with a scaffold N50 0f 19.24 Mb. Manual curation of the scaffolded assembly produced 61 scaffolds spanning 281.14 Mb. Successive purging of this assembly removed 12.14 Mb, and the second contamination screen removed 4 Streptophyta scaffolds totaling 2.52 Mb.

The final assembly was 266.48 Mb in 40 scaffolds, of which 16 are chromosome scale and range from 12.45 to 23.40 Mb (Figure 4). The 16 chromosomes contain 96.9 % (258.20 Mb) of the assembly with the remaining 3.1 % (8.28 Mb) distributed across 24 unplaced scaffolds. This assembly has a scaffold N50 of 15.18 Mb, L50 of 8, and a contig N50 of 4.46 Mb across 115 contigs. Furthermore, the assembly contains 76 gaps totaling 7,600 bp, a GC content of 55.69%, a consensus quality value (QV) of 63.17, and k-mer completeness of 83.33%.

**Figure 4.**
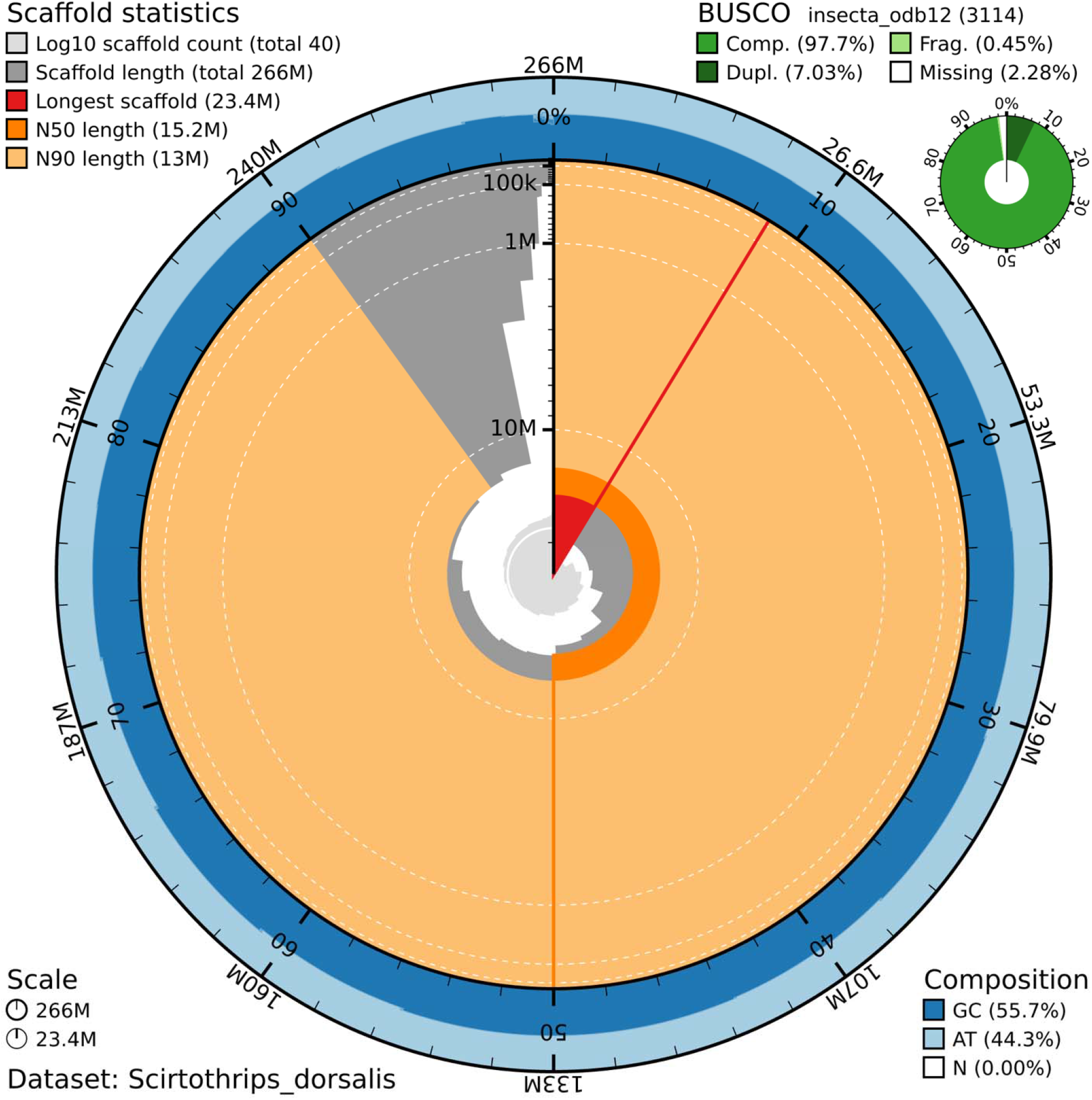
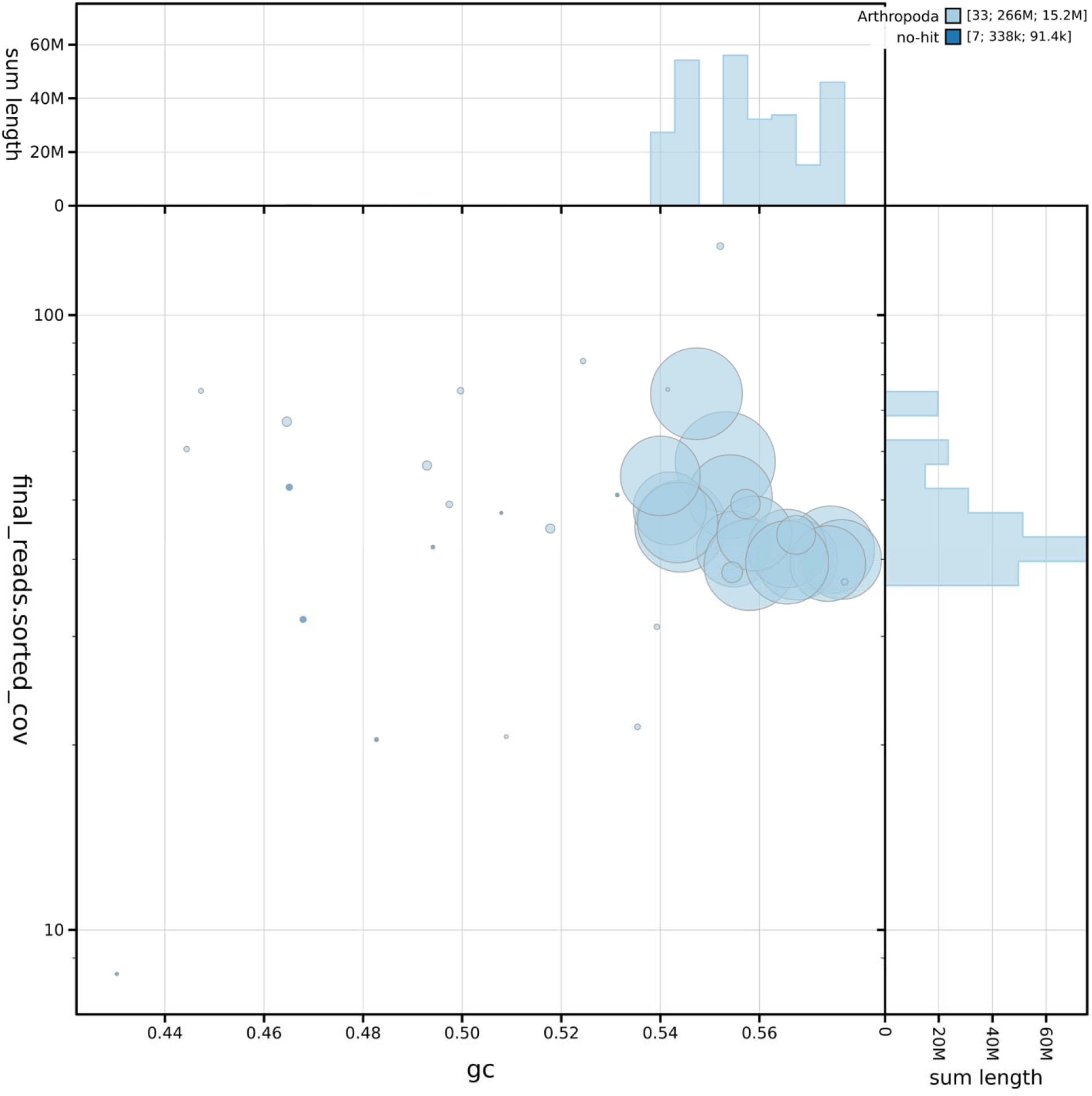
Assembly metrics for the final *Scirtothrips dorsalis* assembly. The first panel is a snail plot of the 266 Mb assembly, which is made up of 40 scaffolds. The grey spiral traces the scaffold length distribution, with the longest scaffold (23.4 Mb) in red, the scaffold N50 (15.2 Mb) in dark orange and the N90 (13.0 Mb) in pale orange. The outer ring shows base composition, with GC content (55.7%) in dark blue and AT content (44.3%) in light blue, and the inset gives BUSCO completeness against the insecta_odb12 set of 3,114 genes, of which 97.7% were complete (7.03% of them duplicated), 0.45% fragmented and 2.28% missing. The second panel is a blob plot of the same assembly, in which each scaffold is a circle placed by its GC proportion and its mapped read coverage, sized by length and colored by phylum-level taxonomic assignment. The 33 scaffolds with a hit sit together in one Arthropoda cluster near a GC proportion of 0.55 and at consistent coverage, and the seven without a hit are all small (338 kb in total), so no contaminant sequence is evident. The histograms along the edges show how total scaffold length is distributed across GC and coverage

BUSCO completeness assessment of the final genome assembly against the Insecta_odb12 dataset showed that the assembly was 97.7% complete, of which 90.7% were single copy and 7.0% duplicated, with 0.4% fragmented and 1.8% missing. It is important to note that duplication greatly reduced at each stage of the assembly, from 46.5% in the initial assembly to 12.3% after the first purge and 7.0% in the final assembly (Figure 5). Scaffolding using Hi-C data changed duplication a little, from 12.3 % to 12.0%, while the second round of purging accounted for most of the remaining reduction in duplication.

**Figure 5.**
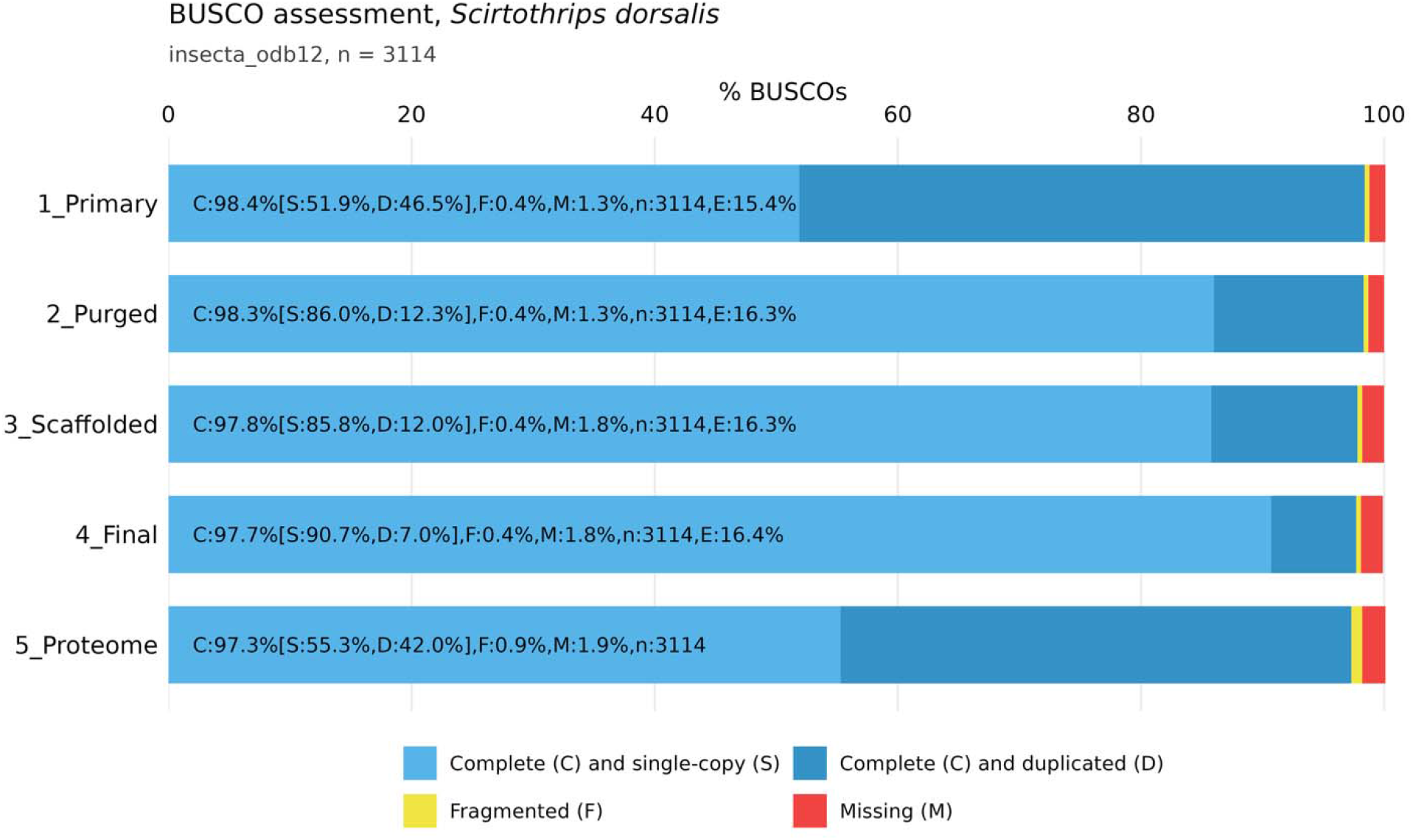
BUSCO completeness against insecta_odb12 at all stages of genome assembly and for the final proteome. The stages are primary assembly using hifiasm, first round of haplotig purging, Hi-C scaffolding and post curation. The proteome was assessed on all predicted transcripts; therefore, duplication corresponds to alternative isoforms.

### Repeat annotation

A total of 455 repeat families were identified by Earl Grey in the *S. dorsalis* genome, of which 439 could not be assigned to a known class, reflecting the limited representation of Thysanoptera in curated repeat databases. Interspersed repeats covered 28.77% of the assembly while 20.30% of the genome was soft masked for gene prediction (Table 3).

**Table 3.** Repeat content of the *Scirtothrips dorsalis* genome.

| Classification | Coverage (bp) | % of genome | Families |
| --- | --- | --- | --- |
| DNA transposons | 2,160,665 | 0.811 | 9 |
| Rolling circle (Helitron) | 2,440,867 | 0.916 | 551 |
| LTR | 485,103 | 0.182 | 6 |
| LINE | 226,235 | 0.085 | 3 |
| Penelope | 37,488 | 0.014 | 1 |
| SINE | 0 | 0.000 | — |
| Unclassified | 56,178,537 | 21.082 | 407 |
| Simple repeat, microsatellite and RNA | 15,123,408 | 5.675 | 1,963 |
| **Total interspersed repeat** | **76,652,303** | **28.765** | — |
| Non-repetitive | 189,825,266 | 71.235 | — |

A large portion of the genome (21.08%) comprised of unclassified elements, whereas the most abundant classified elements were rolling circle transposon followed by DNA transposons, LTR elements and lastly LINEs, while no SINE were recovered. Additionally, 5.68% of the genome consisted of microsatellites, structural RNAs, and simple repeats, which were annotated but not masked. The rolling-circle elements recovered by HELIANO were distributed in 551 families with each family, averaging approximately 4 copies. The most abundant DNA transposons were CMC-EnSpm drawn from 9 families.

### Gene prediction and annotation

The merged annotation from BRAKER4 and GALBA consisted of 20,557 protein coding genes and 29,549 transcripts. Of the total transcripts recovered, 88.9% had their coding sequences split across two or more exons (were multi-exonic), while 91.4% carried a complete coding sequence with both start and stop codons. BUSCO assessment of the annotation against the Insecta_odb12 dataset showed a score of C:97.2% [S:89.8%, D:7.4%], F:0.9%, M:1.9%, which was greater than that of GALBA and RNA-seq-based BRAKER4 (Figure 6)

**Figure 6.**
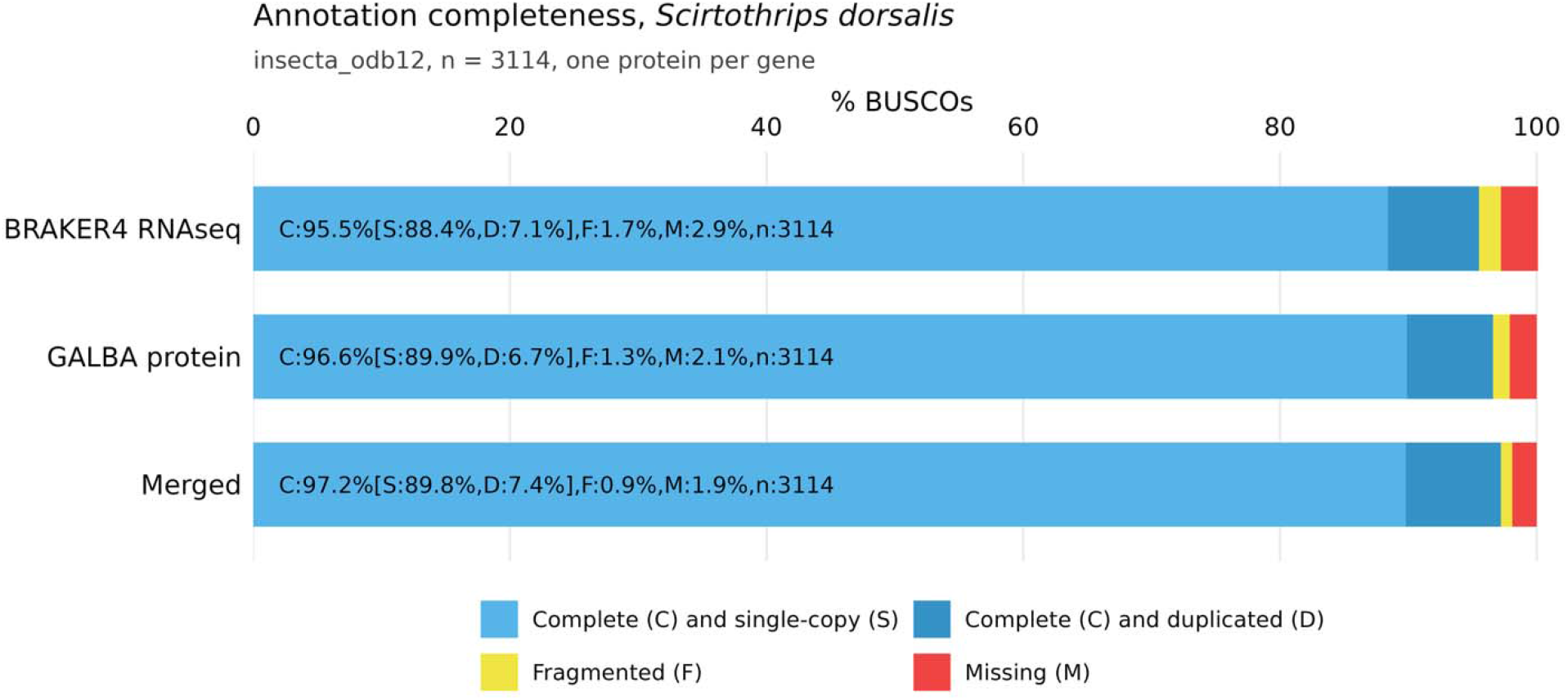
BUSCO completeness of the BRAKER4, GALBA and merged annotations assessed at one protein per gene.

A comparison of the merged annotation against each annotation by BRAKER4 and GALBA revealed that 17,691 genes agreed one-to-one with those of BRAKER4, while 14,515 genes agreed in a similar fashion to those of GALBA. However, of the merged set, 599 had no BRAKER4 counterpart and were only recovered from protein homology alone whereas 4,080 had no GALBA complement and were thus derived from RNA-seq evidence alone. During merging, 170 BRAKER4 and 327 GALBA genes were not retained in the final set.

Functional evidence from at least one source was assigned to 74.0% (15,215 genes) of the annotation. InterPro domains were the most extensively assigned at 14,034 genes, followed by Pfam domines at 12,758 genes, eggNOG orthologous groups at 12,659 genes while 7,605 genes were assigned to KEGG orthologues corresponding to 5,149 distinct KO terms across 379 pathways. The sources complemented each other in that 2,470 genes carried InterPro evidence without eggNOG, 1,095 genes carried the eggNOG in the absence of InterPro, while 9,252 genes carried InterPro, Pfam, eggNOG, GO and COG assignments in combination (Figure 7).

**Figure 7.**
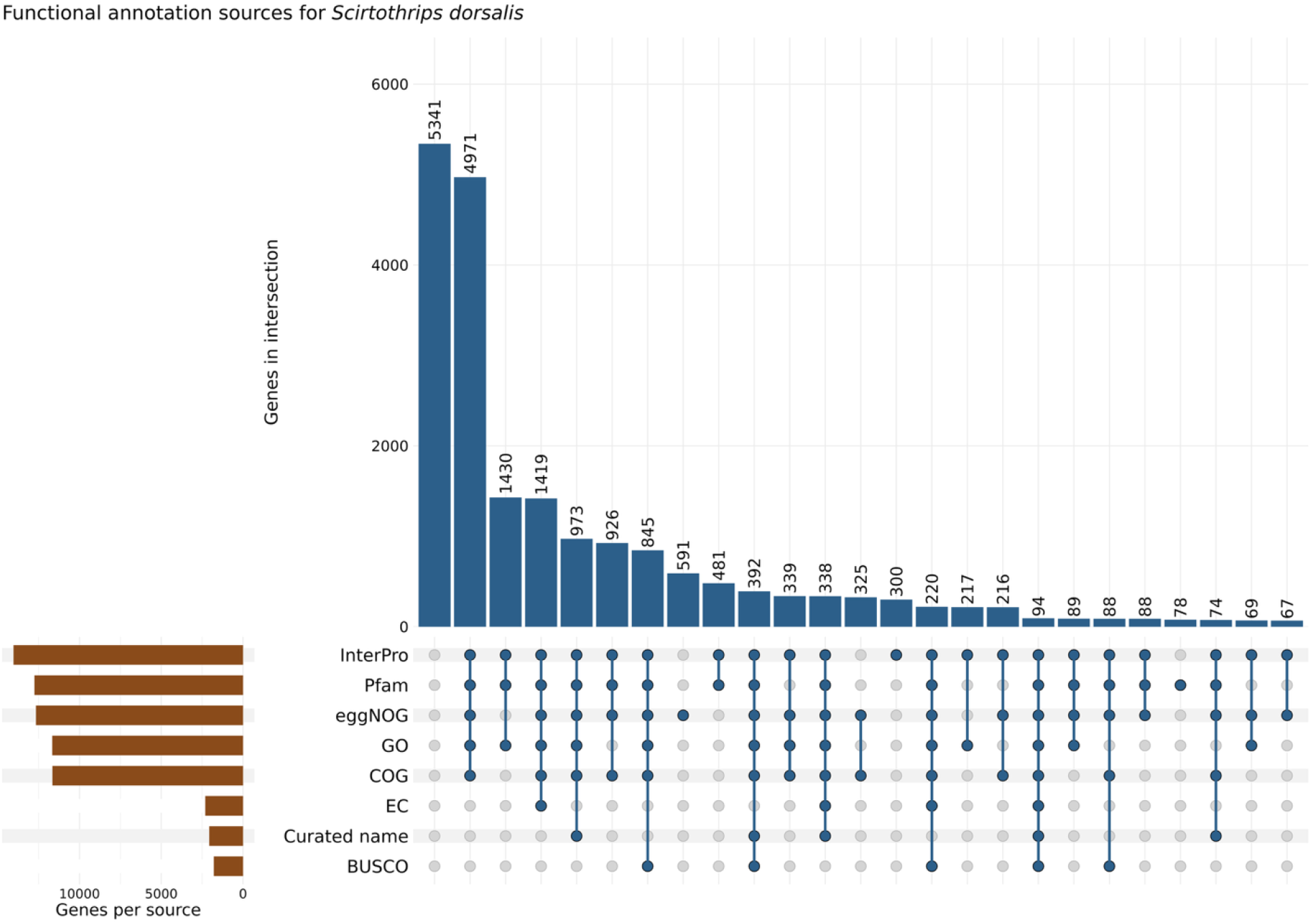
Overlap of functional annotation sources across the 20,557 predicted genes. Bars show the number of genes in each intersection, left-hand bars show the total assigned by each source, while the circles indicate contributions of each source to the intersection

Homology searches against UniProtKB returned hits for 46.8% of transcripts in Swiss-Prot and 80.7% in the arthropod subset of TrEMBL. Non-coding RNA annotation identified 1,077 transfer RNA genes alongside 645 tRNA-derived pseudogenes, 972 ribosomal RNA genes of which 955 were 5S rRNA dispersed across all 16 chromosomes, 173 microRNAs, 134 small nuclear RNAs, 23 other non-coding RNAs, 20 small nucleolar RNAs and 7 signal recognition particle RNAs.

### Chromosomal architecture

Gene density, repeat content, GC content and non-coding RNA density were profiled in 100 kb windows along each chromosome (Figure 8). The repeat landscape was dominated by unclassified elements along the full length of the chromosome, whereas classified repeats were sparse and formed discrete peaks instead of a continuous background.

**Figure 8.**
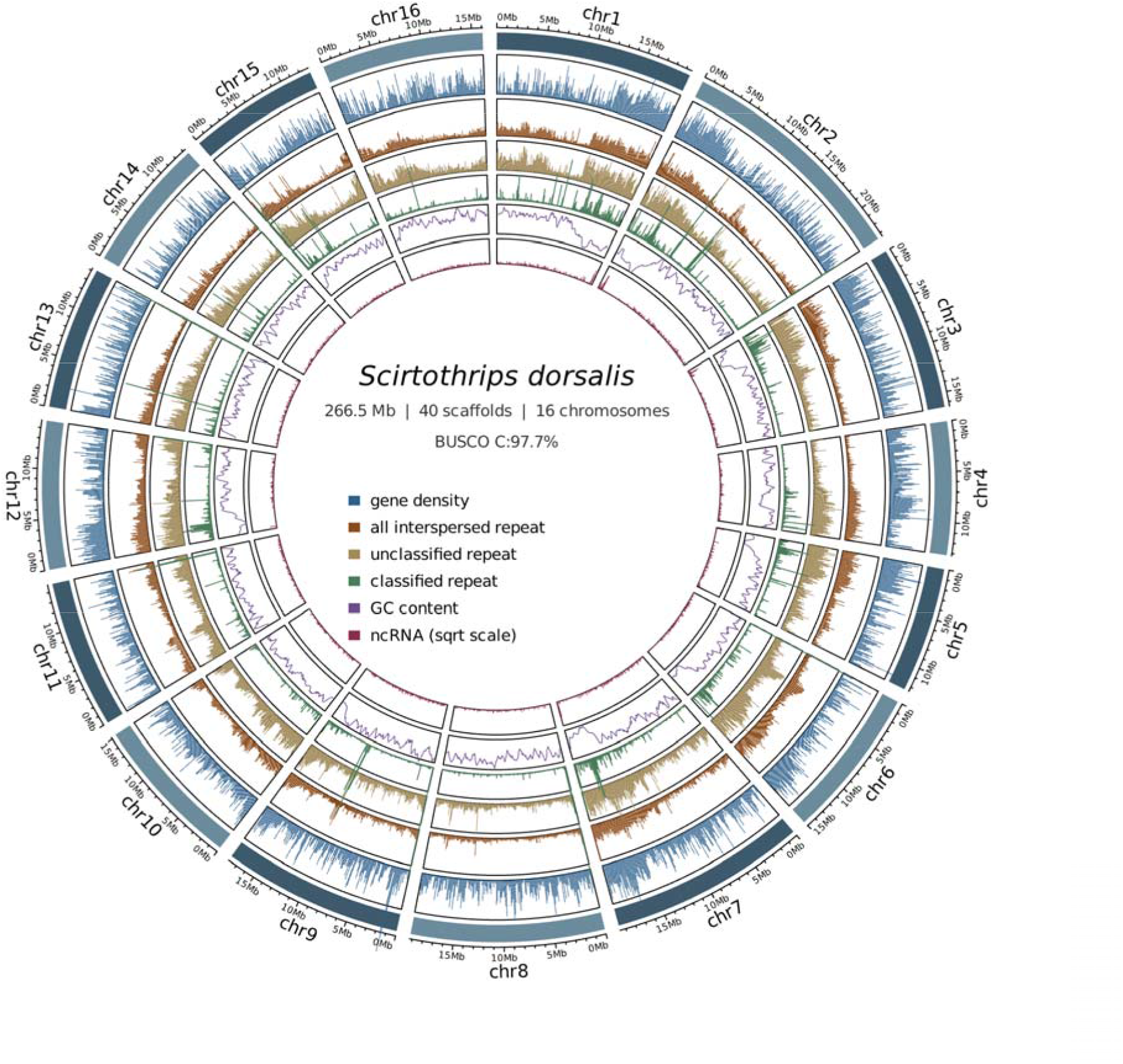
Chromosome-scale architecture of the *Scirtothrips dorsalis* assembly. The outermost ring shows the 16 chromosomes with a 5 Mb scale. Tracks from outside to inside show gene density, all interspersed repeat content, unclassified repeat content, classified repeat content, GC content smoothed over 500 kb, and non-coding RNA density, calculated in 100 kb window along each of the 16 chromosome-scale scaffolds. Non-coding RNA density is plotted on a square root scale, while the 24 unplaced scaffolds, totaling 8.28 Mb, are not shown.

Direct reconstruction of satellite repeats using HiFi reads further resolved the repeat landscape. At the extreme ends of chromosomes 2, 6, 8, 9 and 11, arrays of 5 to 44 kb were formed by a 533 bp unit at relative positions between 0.000 and 0.005 or between 0.996 and 0.999.

At the ends of 5 other chromosomes, terminal arrays of the canonical insect telomeric motif TTAGG (Y. Zhou et al. 2022) were recovered in blocks of 568 to 908 copies on chromosomes 10, 12, 14 and 15, and 28 copies on chromosome 7. Given that no chromosome carried both the satellite and the telomeric arrays, the satellite likely occupies a subtelomeric position from which the assembly did not extend to the telomere. A distinct 1,040 bp unit was found to occupy internal positions on chromosomes 3, 4 and 8 at relative positions 0.515 to 0.878, and a 469 bp unit formed a 38.7 kb array at relative position 0.498 on chromosome 7. At the end of chromosome 7, 5 tandem copies of a 15.3 kb unit were found to be consistent with a ribosomal DNA repeat.

### Phylogenetic placement

The final super matrix contained 3,019 single copy orthologs and 1,610,654 aligned amino acid positions across 13 taxa, 453,374 of which were sparingly informative. Missing data ranged from 3.6% in *T. hawaiiensis* to 25.9% in *T. tabaci* and was 10.2% in *S. dorsalis* and 8.9% in *S. hansoni*. The concatenated and coalescent analyses produced identical topologies (Figure 9).

**Figure 9.**
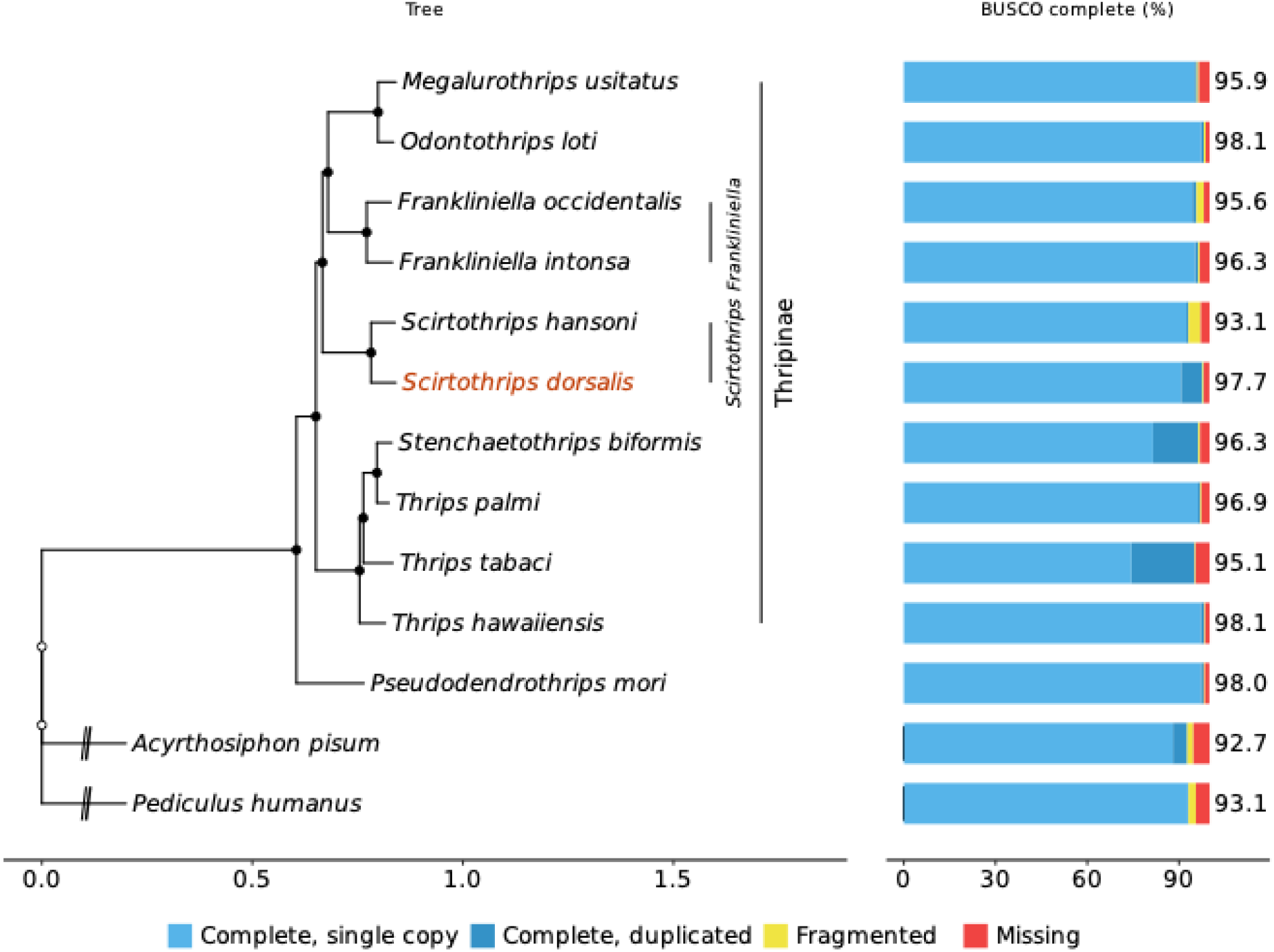
Phylogenetic position of *Scirtothrips dorsalis* (red) among available thrips genomes, based on 3,019 single copy orthologs. The tree is rooted on *Pediculus humanus*, and outgroup branches are shortened for display (//). Filled circles indicate maximum support in the concatenated analysis. Scale bar shows substitutions per site. The right panel shows BUSCO completeness (insecta_odb12) for each species.

Every node in the concatenated tree received 100% SH-aLRT and ultrafast bootstrap support, and all but two nodes in the coalescent tree received local posterior probabilities of 1.00 (0.98 and 0.99).

*Scirtothrips dorsalis* was recovered as a sister group to *S. hansoni*, therefore confirming its placement within the genus. The *Scirtothrips* genus was in turn a sister clade to one containing *Frankliniella* and *Megalurothrips* + *Odontothrips* while this group was sister to the clade consisting of *Thrips* and *Stenchaetothrips biformis*. Together, all Thripinae formed the sister group of Dendrothripinae that contained *Pseudodendrothrips mori*. Within the Thrips clade, *S. biformis* was nested within the genus as a sister to *T. palmi*, rendering *Thrips* paraphyletic. Buckman et al. (2013) also placed *S. biformis* within the *Thrips* genus as a sister to *Thrips nigropilosus*. Hu et al. (2023) also recovered a close relationship between *S. biformis* and *T. palmi* and approximated that they might have diverged about 26 million years ago. However, the analysis by Hu et al. (2023) only included a single *Thrips* species and therefore could not truly test the monophyly of the genus. Since three *Thrips* species were sampled here, the present results suggest that *Stenchaetothrips* may belong within *Thrips*, although broader sampling of both genera and their close relatives will be needed to confirm this.

The branch leading to *S. hansoni* (with 0.057 substitutions per cite) was similar in length to the one leading to *S. dorsalis* (with 0.059 substitutions per cite). This indicated that the highly fragmented *S. hansoni* assembly did not distort its position in the tree. On the other hand, the branches separating the main groups within Thripinae were very short (0.013 and 0.015 substitutions per site) compared with the branch leading to *Scirtothrips* (0.115). They were also short in coalescent units (0.035 and 0.071), suggesting that the main lineages of Thripinae diverged rapidly. Agreement between the two methods under these conditions added confidence to the relationships recovered.

## Conclusion

This study presents the first chromosome-level genome assembly for the genus *Scirtothrips*, from the South Asia 1 cryptic species of *S. dorsalis*. The 266.5 Mb assembly, with 96.9% of the sequence placed in 16 chromosome-scale scaffolds, is comparable in size and gene content to other sequenced Thripidae. Phylogenomic analysis placed *S. dorsalis* with *S. hansoni* and closest to *Frankliniella, Megalurothrips* and *Odontothrips*, and supported earlier evidence that *Stenchaetothrips biformis* falls within Thrips. The mitochondrial genome of the Florida colony is a single circular molecule containing the sequence that forms a separate mini-circle in the published South Asia 1 genome, showing that mitochondrial genome structure differs between populations of the same cryptic species. This genome provides a reference for studying the genetic basis of insecticide resistance and host use in *S. dorsalis*, for resolving relationships within the cryptic species complex, and for developing more targeted approaches to managing this pest.

## Data availability

The *Scirtothrips dorsalis* genome assembly has been deposited at DDBJ/ENA/GenBank under accession JCEILV000000000; the version described in this paper is JCEILV010000000. Raw sequencing reads are available in the Sequence Read Archive under BioProject PRJNA1531588 and BioSample SAMN63248410. These records are held private until publication and can be accessed by reviewers through this <u>link</u>. Additionally, the genome annotation comprising the GFF3 file and the predicted protein sequences, has been deposited in Zenodo. All data will be made publicly available upon publication. The mitochondrial genome was not deposited, as it is nearly identical to the published South Asia 1 sequence (KM349827.1 and KM349828.1).

## Acknowledgements

The authors thank Diem Nguyen for her help throughout the project, particularly in collecting the samples sent for sequencing.

